# Cold Storage and Cryopreservation Methods For Spermatozoa of the sea urchin *Lytechinus pictus*

**DOI:** 10.1101/2023.09.14.557644

**Authors:** Victor D. Vacquier, Amro Hamdoun

## Abstract

Sea urchins have contributed greatly to knowledge of fertilization, embryogenesis and cell biology. However, until now, they have not been a genetic model organism because of the long generation times of commonly used species, and lack of tools for husbandry and genetic manipulation. We recently established *Lytechinus pictus*, as a multigenerational sea urchin model, because of its relatively short generation time of 4-6 months and ease of laboratory culture. To take full advantage of this new multigenerational species, methods are needed to biobank and share mutant *L. pictus* sperm. Here, we describe a new extender based on sperm ion physiology before spawning of sperm into seawater. This extender maintains sperm capable of fertilization for at least 5-10 weeks when stored at 0 °C. We use the extender, and the cryoprotectant dimethyl sulfoxide (DMSO), to cryopreserve sperm of both *L. pictus*, and the widely used sea urchin, *Strongylocentrotus purpuratus*. The simple methods we describe work well for both species, achieving > 90% development and producing larvae that successfully undergo metamorphosis to juvenile sea urchins. Sperm of these two species can be frozen and thawed at least twice and still give rise to larvae that undergo metamorphosis.

**Main Points:**

1. Sperm can maintain fertilizing capacity *ex vivo* for 5-10 weeks when stored at 0°C.
2. When freezing in liquid nitrogen no stepwise addition of cryoprotectant, or stepwise drop in temperature are required.
3. A standard fertilization assay is presented to score cleavage stage sea urchin embryos produced by cryopreserved sperm.
4. Sperm frozen and thawed more than once can produce larvae.

## 1. Introduction

Cryopreservation of animal sperm is important for biobanking genomes of research and agriculturally important animals and for conservation of rare and endangered species. Cryopreservation of sperm is also important to preserve mutant genes for distribution to commercial and research laboratories. Methods for animal sperm cryopreservation involve multiple variables and many methods are restricted to closely related species and can even be species-specific.

Sea urchins are members of Phylum Echinodermata, a major animal taxonomic group, which to our knowledge has not utilized mutations and multigenerational studies to discover the function of specific genes. The deficit in methodology is because most sea urchin species have generation times greater than 1 year. Because of CRISPR-Cas9, the creation of transgenic sea urchins is now in progress in several laboratories (21,22,29,42). One California species, *Lytechinus pictus*, is the most promising urchin for multigenerational studies due to its generation time of 4-6 months and ease of laboratory culture (14,24,25). The genome of *L*.*pictus* is sequenced (41) and F2 and F3 generation adults have been created with the multidrug transporter ABCB1 deleted (33,40).

The history of cryopreservation of sea urchin sperm began in 1973 (9), and at present, sperm of about 15 species have been cryopreserved as recently reviewed (11,26-28,31). Cryopreservation of sperm of all animals involves dilution of concentrated semen into an extender solution and then mixing this cell suspension with a cryoprotectant such as dimethyl sulfoxide (DMSO). After various times of incubation, this mixture is frozen in liquid nitrogen (LN), either with or without steps of different temperatures held for various times. When the desired time of storage in LN is reached, thawing is generally as rapid as possible.

Most studies of ion physiology of sea urchin sperm have utilized *S. purpuratus* and a wealth of information exists. Ion channel activities of these sperm regulate: ATP levels, respiration, motility, chemotaxis to egg peptides and induction of the acrosome reaction (3,5-8,12,16,19,23,30). Summarizing the findings of these citations, when in the testis, or when stored as undiluted semen, sea urchin sperm have an internal pH (pHi) of 6.8-7.1, ATP concentrations are highest and the cells are nonmotile. When sea urchin semen is diluted into seawater a coupled exchange of 1 Na^+^ in, to 1 H^+^ out, occurs in seconds and intracellular pH increases to 7.6. The pHi increase activates axonemal dynein ATPase, ATP is hydrolyzed and motility begins. The H^+^ created by dynein activity is effluxed by the Na^+^/H^+^ exchanger. Na^+^ concentration is maintained by the Na^+^/K^+^ ATPase and the intracellular K^+^ concentration is controlled by passive leak through the plasma membrane. A further increase in pHi from 7.6 to 7.8 occurs when sperm undergo the acrosome reaction. The pre-spawn pHi of 6.8-7.1, high ATP and nonmotility can be experimentally maintained in *S. purpuratus* sperm if concentrated semen is diluted into seawater buffered with 5-10 mM HEPES pH 7.0 and 40 mM extra KCl (final K^+^ = 50 mM).

## 2. Material and Methods

### 2.1 Biological Material

Adults of both species were collected at La Jolla, California, under a scientific collecting permit to our institution issued by California Department of Fish and Game. Adults were maintained in aquaria circulating with the ocean at controlled temperatures of 12°C *(S.purpuratus)* and 20°C *(L. pictus)* and fed the brown kelp, *Macrocystis pyrifera*. Adults were induced to spawn by injection of 0.55 M KCl. Undiluted semen was stored in tubes in ice referred to as “dry sperm”. Eggs were stored at 15°C (*S. purpuratus*), or 22°C (*L.pictus*) for no longer than 12 hours.

### 2.2 Long-term viability of sperm stored at 0°C

Before dilution into filtered seawater (FSW), undiluted semen of these two species has a fairly constant sperm cell number of 4 (*S. purpuratus*) and 3 (*L. pictus*) x 10^10^ cells per ml (34,36). Blunt micropipette tips used in all experiments had openings of 1-2 mm to avoid shear force injury to cells. The sperm extender, designated KH7, is composed of FSW (filtered seawater) containing an extra 40 mM KCl, 5 mM HEPES buffer adjusted to pH 7.0 with 1N NaOH and 1N HCl. In all experiments, undiluted semen was diluted 1 to 100 v/v in KH7 or FSW, gently mixed and the tubes packed in ice. In the first part of this study, semen was diluted 1 to 100 in KH7 or FSW, and packed in ice in 15 ml screw-cap tubes that were placed in a styrofoam box, which was placed in a 4°C cold room and the sperm tested occasionally for fertilization ability. The sperm storage temperature never rose above 0°C as determined by mercury and alcohol thermometers.

### 2.3 Cryopreservation in liquid nitrogen

The standard method to cryopreserve sperm was as follows. Pure DMSO (Fisher Scientific) was distributed in nine 12 ml glass conical centrifuge tubes in volumes of 60, 90, 120, 150, 180, 210, 240, 270, 300 μl and the tubes kept at room temperature (22C). Different volumes of sperm, diluted 1:100 in KH7 or FSW, were pipetted into each DMSO containing tube and rapidly mixed to make a final volume of 1 ml. For example, 940 μl of sperm in KH7 or FSW were rapidly mixed with 60 μl DMSO to make 1 ml with 6% v/v DMSO (0.84 M DMSO). Likewise, 700 μl of sperm in KH7 or FSW was rapidly mixed with 300 μl DMSO to make 1 ml with 30% DMSO (4.26 M).

The nine 12 ml tubes of sperm in various DMSO concentrations were packed in ice for 60 min with gentle mixing by swirling the tubes at 10 min intervals. This equilibration time could be extended 4 hours without decreasing fertilization results. The 1 ml in each tube was then divided into two portions of 500 μl and transferred to two 2 ml capacity cryovials. The 18 cryovials were tightly capped and placed in a 5 cm square cardboard cryobox that was immediately placed in a LN dewar (VWR CryoPro).

### 2.4 Thawing cryovials

Cryoboxes were kept in LN for 1 to 152 days. To thaw cryovials, the cryovials were dropped into 1.5 liters of tap water at 15°C *(S.purpuratus)*, or 20°C (*L.pictus)* and the water stirred vigorously by hand for 3 min. The completely thawed cryovials were then removed from the water and liquid in the tube cap brought down by hitting the vial bottom on the bench top. The tubes were then placed on ice until used.

### 2.5 Fertilization test used in all experiments

Six-well flat bottom culture plates (Olympus Plastics Cat# 25-100, 16 ml capacity) were used for all fertilization tests. New plates were cleaned by exposure to 2 ml 1 N NaOH for 12 hours, followed by 2 ml 1 N HCl for 12 hours. The plates were then soaked in deionized water, 12 hours, rinsed and dried. After the first use the plates were simply washed in tap water and air dried. For the fertilization test, the plates were put on a rotary table (80 rpm) with 2 ml FSW containing 1 mM HEPES, pH 8.1 (FSW/HEPES).

Spawned eggs (0.1-0.4 ml settled volume) were resuspended in 45 ml FSW and slowly poured twice through nylon netting of 85 μm pore size for *S. purpuratus*, (egg diameter 80 μm), or 120 μm for *L. pictus* (egg diameter 111 μm) to remove debris and loosen the egg jelly coat. The egg suspension was then poured into a 50 ml graduated conical plastic tube and hand centrifuged slowly for 40 revolutions of the hand crank to sediment the eggs. The supernatant was completely aspirated away and the egg pellet resuspended in 30 ml FSW/HEPES pH 8.1, the tube capped and inverted slowly 30 times, and the eggs kept in suspension while twelve 10 μl samples were removed, placed on a clear plastic dish and the eggs counted using a dissecting microscope. Egg concentration was further diluted in FSW/HEPES until 15-30 eggs were found in a 10 μl drop. The volume of egg suspension (100-200 μl) containing between 100-300 eggs was determined and added to each well of the 2 ml FSW/HEPES rotating at 80 rpm to keep the eggs from settling on and sticking to the well surface.

Sperm, which had been stored for for various times at 0°C, or frozen in LN and thawed, were then added to the rotating plates in volumes of 3 to 100 μl per well and 90 rpm rotation continued until the first cleavage division (*L. pictus* ∼70 min at 22°C, *S.purpuratus* ∼110 min at 15°C). The greater than 20-fold dilution of the DMSO in the sperm did not affect fertilization. Rotation was then slowed to 60 rpm during the cleavage period. Adding 100 μl of sperm suspension to 2 ml of FSW/HEPES decreased the pH of the fertilization mixture from 8.1 to 7.8.

Sperm concentrations in the fertilization mixture were determined by removing 40 μl from each inseminated well and counting the sperm cells at 400 x magnification using a hemocytometer and phase contrast microscope. In most experiments the fertilization ability of eggs was determined by inseminating an egg sample in FSW/HEPES with non-frozen sperm from another male.

### 2.6. Scoring percent development

Embryogenesis in *L. pictus* is more than twice as fast as in *S.purpuratus* (14,24,25). Percent development was determined by photographing each well of the 6-well plates between the 4-cell cleavage stage and blastula stage. Plates were rotated by hand to gather the eggs and embryos in the center of the well prior to photographing the well. Photographs were printed on 21 x 27 cm paper and the number of unfertilized eggs and embryos counted by hand to determine percent development. We did not score percent fertilization by counting eggs with elevated fertilization envelopes. Envelope elevation can be misleading due to artificial activation caused by physical damage to eggs. To stop blastulae swimming during photography, 25 μl of 10% glutaraldehyde in SW was added to each well. If long term development was planned, 25 to 50 blastulae were removed by hand pipetting and recultured in 8 ml FSW in 5 cm diameter petri plates before the remainder of the dish was fixed in glutaraldehyde. Images were acquired with a Leica M165FC compound microscope with a 1X Plan Apo objective and a Canon EOS 60D camera.

### 2.7 Long term culture of embryos and larvae

Swimming blastulae were picked out by hand pipetting and transferred to 8 ml FSW in 5 cm diameter petri plates that were incubated at 15°C for *S.purpuratus* and 22°C for *L. pictus*. At gut completion the embryos were fed the unicellular alga *Rhodomonas lens* and were cultured as previously described (14,24,25,38). Complete changes of the FSW occurred every 5-7 days. The

*L. pictus* larvae can undergo metamorphosis in 17-21 days, whereas *S. purpuratus* require at least twice that time. All data are presented in table form as percent development with the number rounded to the nearest whole number.

## 3. Results

### 3.1 Fertilization ability after sperm storage at 0°C in KH7 and FSW

*L. pictus* sperm diluted 1:100 in KH7 and kept packed in ice can fertilize eggs for 32 days, whereas if stored as undiluted semen (“dry”) and then diluted 1:100 in KH7 15 min before insemination, almost all their fertilizing ability is lost by 21 days. The *L.pictus* sperm diluted 1:100 in FSW retain no fertilization ability after 32 days at 0°C. (Table 1).

**Table 1.** Percent embryonic development resulting from storage at 0°C of *Lytechinus pictus* sperm three different ways. First, semen (“dry sperm”) was diluted 1:100 in KH7 buffer (filtered seawater with 5 mM HEPES and 40 mM extra KCl, pH 7.0). Second, semen was diluted 1:100 in filtered seawater (FSW). Third, semen was left undiluted “dry”. All three ways used 15 ml screw cap tubes for storage. The undiluted semen was diluted 1:100 in FSW 15 min before insemination. Fertilization tests were performed after 21, 32, 39 and 46 days of storage at 0°C. The percentage fertilization decreased with time, but KH7 buffer retained fertilization ability for one month. This table represents scoring 8,375 eggs and embryos. After insemination in the 6-well plate with 100 μl sperm, the sperm concentration in the total volume of 2.1 ml was approximately 1.4 x 10^7^ sperm per ml.

|  | 21d |  |  | 32d |  | 39d | 46d |
| --- | --- | --- | --- | --- | --- | --- | --- |
| *ul sperm* | KH7 | FSW | Dry | KH7 | FSW | KH7 | KH7 |
| 100 | 99 | 4 | 6 | 76 | 0 | 12 | 4 |
| 50 | 100 | 4 | 3 | 51 | 0 | 7 | 2 |
| 25 | 100 | 0 | 3 | 36 | 0 | 7 | 2 |
| 12 | 95 | 0 | 2 | 30 | 0 | 5 | 8 |
| 6 | 67 | 0 | 1 | 12 | 0 | 3 | 0 |
| 3 | 59 | 0 | 1 | 8 | 0 | 4 | - |

The *S. purpuratus* sperm diluted in KH7 or FSW and stored 53 days at 0°C show that FSW is better than KH7 for long term storage in this species (Table 2). The *S. purpuratus* sperm stored “dry” as undiluted semen at 0°C for 20 days had no fertilizing ability. Seventy days at 0°C is the longest we have stored *S. purpuratus* sperm and kept them highly fertilizable (not shown). For sperm of both species, the viability durations at 0°C are long enough to allow shipment of sperm without freezing in LN.

**Table 2.** Percent embryonic development resulting from storage at 0°C of *Strongylocentrotus purpuratus* sperm kept undiluted (dry) or diluted 1:100 in either FSW, or KH7. At 53 days KH7 buffer is not as protective as FSW. The table represents scoring 8,993 eggs and embryos. In the fertilization mixture of 2.1 ml, in the 100 μl sperm well, the sperm concentration was approximately 1.9 x 10^7^ sperm/ml.

|  | 20d |  |  | 41d |  | 53d |  |
| --- | --- | --- | --- | --- | --- | --- | --- |
| *ul sperm* | KH7 | FSW | Dry | KH7 | FSW | KH7 | FSW |
| 100 | 99 | 100 | 0 | 87 | 90 | 97 | 97 |
| 50 | 100 | 100 | 0 | 96 | 91 | 85 | 93 |
| 25 | 99 | 100 | 0 | 96 | 92 | 60 | 95 |
| 12 | 100 | 98 | 0 | 94 | 92 | 41 | 96 |
| 6 | 99 | 98 | 0 | 96 | 95 | 34 | 96 |
| 3 | 99 | 98 | 0 | 96 | 98 | 23 | 95 |

### 3.2 Lytechinus pictus sperm short duration freezing in liquid nitrogen

DMSO at 24% yielded few embryos, and 27% and 30% yielded no embryos in both species. Table 3A presents data from one experiment run in quadruplicate using four 6-well plates (Plates A-D) and sperm and eggs from one male and one female. These sperm were frozen in liquid nitrogen for 20 days, thawed, and 100 μl used to inseminate each well and the unused sperm immediately refrozen. Each well had 100-300 eggs in 2 ml FSW/HEPES. With the exception of the 21% DMSO wells, replicates of the other concentrations of DMSO are in good agreement as shown by the reasonable standard deviations.

**Tables 3A & 3B.** Percent development using *Lytechinus pictus* sperm short duration freezes in liquid nitrogen. **A. Freeze 1**. *Lytechinus pictus* sperm were prepared for cryopreservation as described, DMSO concentrations ranged from 6%-21%. At 20 days the cryovials were thawed and fertilization tests done in quadruplicate using four 6-well plates (A-D). Each well of 100-300 eggs was inseminated with 100 μl of sperm (average sperm concentration in the fertilization mixture was 8.1 x 10^6^/ml. Residual sperm was immediately refrozen. Photomicrographs were taken at 24 hours. **B. Freeze 2**. After nine additional days in LN the cryovials were thawed and the second fertilization test done. DMSO concentrations from 9% to 18% were protective for *Lytechinus pictus* sperm. SD = standard deviation.

|  | Plate |  |  |  |  |  |
| --- | --- | --- | --- | --- | --- | --- |
| % DMSO | A | B | C | D | ave | SD |
| 6 | 87 | 92 | 91 | 81 | 88 | 5.0 |
| 9 | 90 | 94 | 84 | 89 | 89 | 4.1 |
| 12 | 93 | 93 | 94 | 93 | 93 | 0.6 |
| 15 | 91 | 93 | 86 | 81 | 88 | 5.4 |
| 18 | 83 | 80 | 74 | 80 | 79 | 3.8 |
| 21 | 59 | 41 | 24 | 37 | 40 | 14.5 |

|  | Plate |  |  |  |  |  |
| --- | --- | --- | --- | --- | --- | --- |
| % DMSO | A | B | C | D | ave | SD |
| 6 | 31 | - | 30 | 17 | 26 | 7.8 |
| 9 | 76 | 82 | 80 | 88 | 82 | 5.0 |
| 12 | 85 | 91 | 90 | 95 | 90 | 4.1 |
| 15 | 92 | 86 | 93 | 49 | 80 | 20.9 |
| 18 | 85 | 83 | 92 | 95 | 89 | 5.7 |
| 21 | 28 | 24 | 71 | 46 | 42 | 21.4 |

The sperm used in Table 3A were refrozen in LN for 9 days and the fertilization test run again in quadruplicate to produce the data shown in Table 3B. Comparing the averages of the 4 wells in A and B shows that the second freeze/thaw cycle was detrimental to sperm in 6% DMSO, but the four well averages for 9%-18% DMSO show good protection of sperm, except for the one outlier of 49% in 15% DMSO (Plate D). The standard deviations are low for 6%, 9%, 12% and 18% DMSO showing that most of the time the four identical wells agree favorably. These data show that *L.pictus* sperm can tolerate two freeze/thaw cycles under our conditions and still yield high percentages of embryos.

### 3.3 Lytechinus pictus sperm long duration freezing in liquid nitrogen

Sperm from one *L. pictus* male was prepared for cryopreservation, divided into duplicate sets of cryovials, and one set frozen for 50 days (Table 4A Plates A & B) and the second set for 114 days (Plates C & D). Eggs from two females had to be used and 100 μl sperm inseminated each well of eggs. The data in Table 4A show that in 6%, 9% and 12% DMSO the 50 day and 114 day wells are fairly comparable between the two freeze durations, with low standard deviations among the four plates. The 15% 18%, 21% and 24% DMSO wells yielded higher percent development in the 50 day wells as compared to 114 day wells (Table 4A). The lower percent development in the 114 day sperm could be due to the extended time in LN, or because the eggs came from different females.

**Tables 4A & 4B.** *Lytechinus pictus* longer duration freeze. **A. Freeze 1**. DMSO concentrations ranged from 6%-24% v/v and duplicate plates were used at each concentration. Sperm from one male was prepared for cryopreservation and divided into two cryovials for each DMSO concentration and frozen 50 days (columns A & B), or 114 days (C & D) before testing. Wells were inseminated with 100 μl and cryovials refrozen. Photomicrographs were taken in 24h. **B. Freeze 2**. The 50 days F1 sperm were refrozen for 83 days (total time in LN =133 days) and tested for fertilization (A’ & B’). The 114d sperm were refrozen for 38 days and fertilization tested (C’ & D’). Data in this table are from scoring 8,513 unfertilized eggs and embryos.

|  | F1=50d |  | F1=114 |  |  |  |
| --- | --- | --- | --- | --- | --- | --- |
| % DMSO | A | B | C | D | ave | SD |
| 6 | 87 | 92 | 85 | 85 | 87 | 3.1 |
| 9 | 89 | 89 | 86 | 86 | 88 | 1.4 |
| 12 | 86 | 87 | 85 | - | 86 | 1.0 |
| 15 | 92 | 91 | 47 | 76 | 77 | 20.9 |
| 18 | 92 | 94 | 71 | 80 | 84 | 10.8 |
| 21 | 90 | 86 | 69 | 63 | 77 | 13.0 |
| 24 | 33 | 42 | 21 | 25 | 30 | 12.9 |

|  | F2=83d |  | F2=38d |  |  |  |
| --- | --- | --- | --- | --- | --- | --- |
| % DMSO | A' | B' | C' | D' | ave | SD |
| 6 | 25 | 17 | 4 | 3 | 12 | 10.6 |
| 9 | 40 | 48 | 51 | 41 | 45 | 5.4 |
| 12 | 69 | 59 | - | 79 | 69 | 10.0 |
| 15 | 66 | 70 | 84 | 69 | 72 | 8.0 |
| 18 | 55 | 59 | 48 | 66 | 57 | 7.5 |
| 21 | 42 | 38 | 30 | 39 | 37 | 5.1 |
| 24 | 13 | 11 | 0 | 0 | 6 | 5.0 |

**Table 5.** Percent development of *Strongylocentrotus purpuratus* after cryopreservation of sperm. Sperm from one male was cryopreserved in duplicate cryovials (Freeze 1). Duplicate 6-well plates were run for each DMSO concentration. Half the cryovials were thawed after 37 days and 100 μl of sperm suspension removed and tested (A & B). These vials were refrozen (Freeze 2) 97 days and tested again (C & D). The other half of the vials (Freeze 1) were thawed for the first time at 89 days and tested (E & F). These vials were refrozen (Freeze 2) and thawed for the second time at 31 days and tested. Loss of fertilization potential from the 2^nd^ freeze/thaw cycle is apparent. Eggs from four females were used for this experiment. Control on egg quality was determined by fertilization with nonfrozen sperm and varied from 91-97% fertilization. 7,854 eggs and embryos were scored for these data. Sperm concentration in the fertilization mixture was approximately 8.3 x 10^6^/ml.

| % DMSO | A | B | C | D | E | F | G | H |
| --- | --- | --- | --- | --- | --- | --- | --- | --- |
| 6 | 89 | 87 | 21 | 27 | 94 | 93 | 17 | 16 |
| 9 | 96 | 88 | 40 | 33 | 94 | 91 | 13 | 13 |
| 12 | 89 | 95 | 29 | 26 | 86 | 92 | 20 | 24 |
| 15 | 87 | 73 | 19 | 28 | 74 | 75 | 43 | 42 |
| 18 | 88 | 92 | 62 | 15 | 91 | 91 | 67 | 69 |
| 21 | 95 | 88 | 35 | 18 | 86 | 68 | 43 | 42 |
| 24 | 70 | 50 | 4 | 8 | 24 | 0 | 4 | 7 |

The sperm used in columns A & B were refrozen for 83 days (total time in LN = 133 days) and tested for developmental potential (Table 4B, A’ & B’). The second freeze/thaw cycle yielded lower percentages (4B) than the first cycle (4A), the greatest variation being at both low and high ends of the DMSO concentrations. The second freeze for the sperm used in C & D was for 38 days (C’ & D’) (total time in LN = 152 days). Comparing the averages in 4A to those in 4B suggests that the greatest cryoprotection occurs in 12%, 15% and 18% DMSO. Both short and long duration freezes yield sperm that produce plutei. These data again show that *L. pictus* sperm can tolerate two cycles of freeze/thaw and still fertilize eggs and produce embryos.

### 3.4 Cryopreservation of Strongylocentrotus purpuratus sperm in liquid nitrogen

Sperm from one *S. purpuratus* male was cryopreserved and cryotubes frozen in duplicate. Half the tubes were frozen for the first time for 37 days, thawed and tested in the standard fertilization assay in duplicate plates (Plates A & B). The unused sperm were immediately refrozen for 97 days (total days in LN = 134) and tested again for fertilization potential (Plates C & D). A decrease in fertilization ability occurred with the second freeze/thaw cycle. Except for 18% DMSO in Plate C of 62% the numbers show the same trend in decrease. The second set of cryotubes was frozen for the first time for 89 days and tested (Plates E & F), refrozen for 31 days and tested again (Plates G & H). Decreases in developmental potential were again seen after the second freeze/thaw cycle. The most protective DMSO concentrations were 15% and 18%.

### 3.5 Post-metamorphic juvenile sea urchins of these two species

Cryopreserved sperm were used to fertilize eggs, which developed into larvae with adult rudiments and underwent metamorphosis to become juvenile sea urchins as previously shown for other species using cryopreserved sperm, but not until now shown for these two species. Figure 1 shows seven juvenile *L. pictus* raised for 46 days from eggs fertilized with cryopreserved sperm frozen for 50 days in 15% DMSO. Metamorphosis occurred between 23-28 days post-insemination (5-7). Figure 2 shows three *S.purpuratus* juveniles, six days post-metamorphosis and 53 days post-insemination, the sperm having been cryoprotected in 15% DMSO for 41 days.

**Fig. 1.**
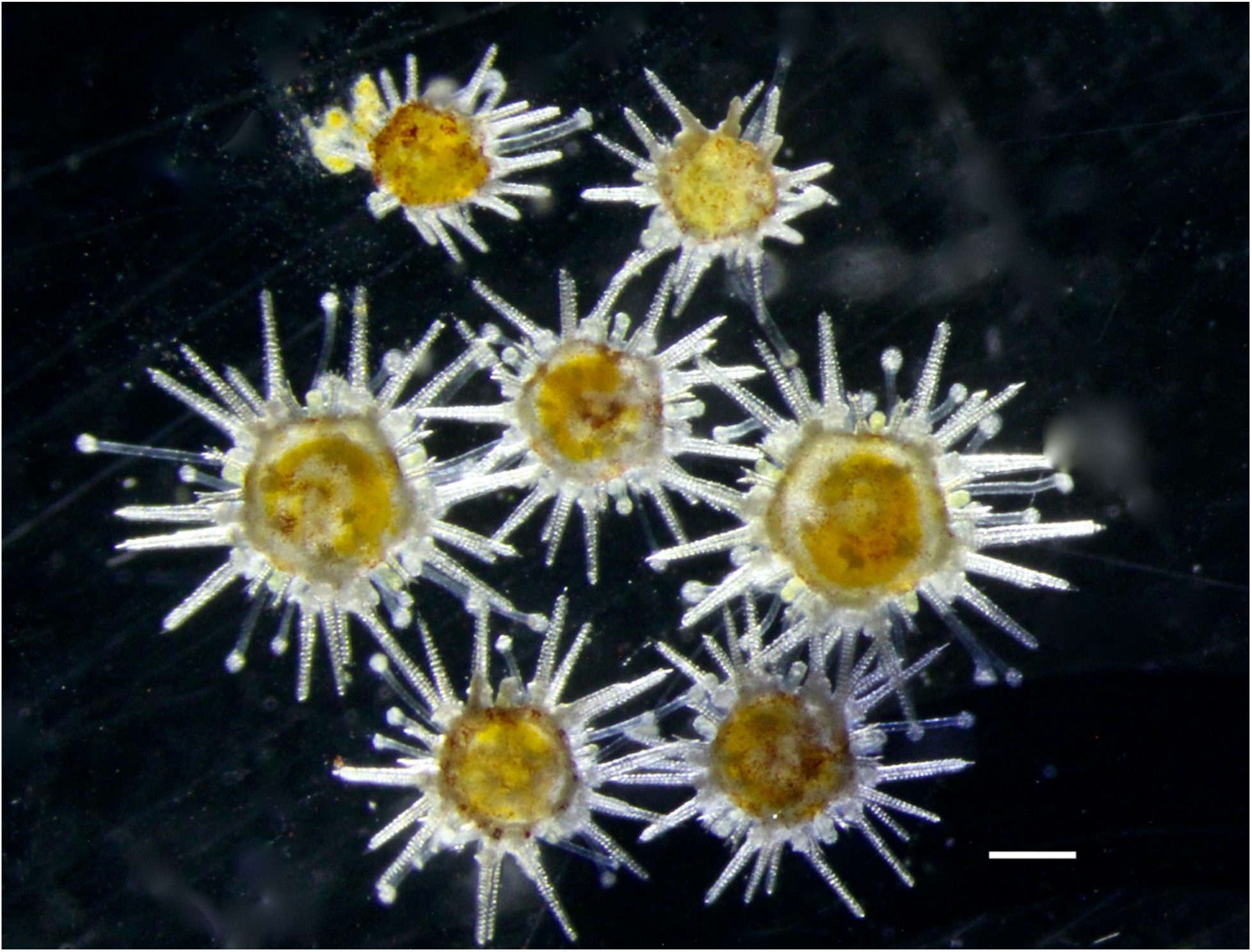
*Lytechinus pictus* post metamorphic juveniles 46 days post insemination. Metamorphosis was 23-28 days post insemination. The sperm were cryopreserved in 15% DMSO for 50 days. Scale bar is 0.5 mm.

**Fig. 2.**
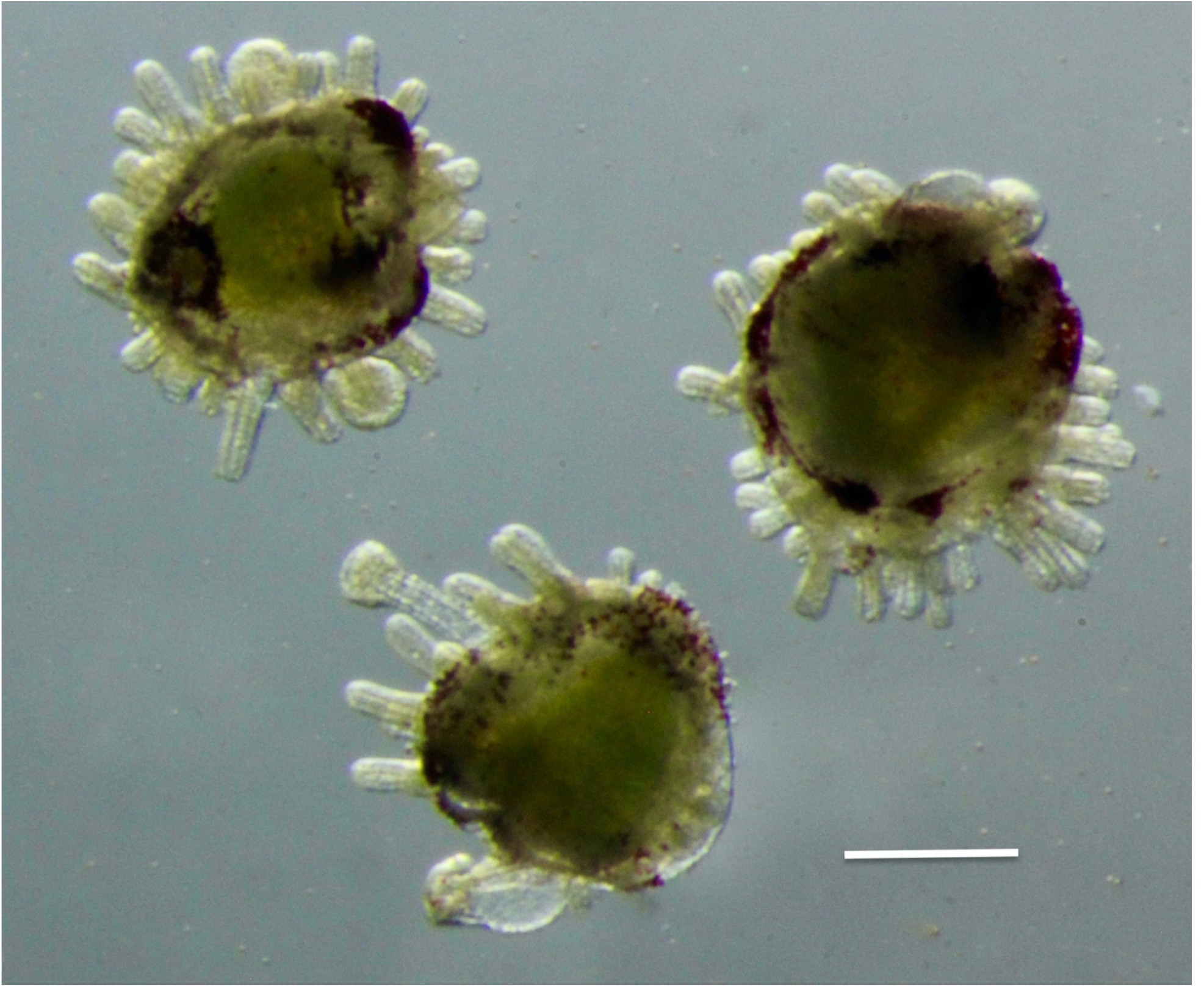
*Strongylocentrotus purpuratus* juveniles 53 days post insemination and six days post metamorphosis. Sperm was cryopreserved 41 days in 15% DMSO. Scale bar is 1 mm.

## 4. Discussion

### 4.1 Extending the fertilizing lifetime of sea urchin sperm by storage at 0°C

The fact that sea urchin sperm fertilizing viability can be extended by dilution in FSW containing 5 mM HEPES (pH 7.0) and an extra 40 mM KCl has been known for years for *S.purp*uratus. However, long term sperm storage has never been carried out at 0°C for the length of reported here for these two species. Routinely keeping *L. pictus* sperm alive for 32 days and *S.purpuratus* sperm for 53 days (70 days maximum) is indeed remarkable. Our results on *S. purpuratus* sperm cold storage agree with results reported for the Japanese sea urchin *Hemicentrotus pulcherrimus*, which showed that 50 mM KCl and 5 mM MES buffer at pH 6 did not increase sperm lifespan over that of seawater (17,18). *H. pulcherrimus* sperm could remain viable for 60 days if the sperm were washed twice in seawater containing antibiotics. However, our data in Table 1 show that with *L.pictus*, 50 mM KCl and pH 7.0 are better at preserving sperm viability than is seawater. Once again this shows that there are significant differences between gametes of sea urchin species.

If these results of long term storage in wet ice at 0°C holds true for other marine invertebrate species, this could be important for sample collection away from the laboratory, because LN dewars would not have to be taken on collecting trips.

### 4.2 Interspecies differences

When *S. purpuratus* sperm are diluted into FSW the Na^+^/H^+^ exchanger activates, and pHi raises from ∼7.1 to 7.6, and dynein ATPase becomes active as does motility. When these sperm are diluted into a pH 7 buffered FSW with 40 mM extra KCl their pHi remains at approximately 7, which is below the sharp pH activation curve of axonemal dynein ATPase (5-7) and the sperm remain immotile.

Sperm of *L. pictus* are remarkably different from *S. purpuratus* because after a few minutes of immotility in KH7 they become motile again. The difference between these two species is that in *S.purpuratus* the acid release is thought to be primarily in the form of protons, whereas in *L. pictus* 57% of the acid is nonvolatile and thought to be protons and 43% is volatile and thought to be CO_2_ (16). Acid release from sperm upon dilution into seawater is easily measured. Figure 3 demonstrates this phenomenon when undiluted semen of these two species is diluted into FSW and the external pH measured at 1 min intervals. In both species there is a rapid acid release for the first six minutes. The sharp drop with *L. pictus* sperm is followed by a steady external pH increase as the volatile acid component leaves the FSW and the external pH reaches 7.6. In *S.purpuratus*, after the sharp drop during the first five minutes, the pH of the external medium remains 7.2-7.3. This low external pH causes the pHi to become acidic and inactivates the dynein ATPase. The molecular phylogeny of seven species of *Strongylocentrotus* is known (20) and it would be interesting to learn if our *S. purpuratus* sperm longevity durations is demonstrable in other *Stongylocentrotus* species. It is interesting that different systems to control sea urchin sperm pHi have evolved in roughly 30-40 million years of divergence (32). The difference in fertilizable lifetime of sperm of these two species when stored at 0°C could be related to this interspecies difference in acid release and pHi. Antibiotics were not used in our long term experiments and even after weeks at 0°C, phase contrast microscopy at 400 X magnification did not show bacterial colonies.

**Fig. 3.**
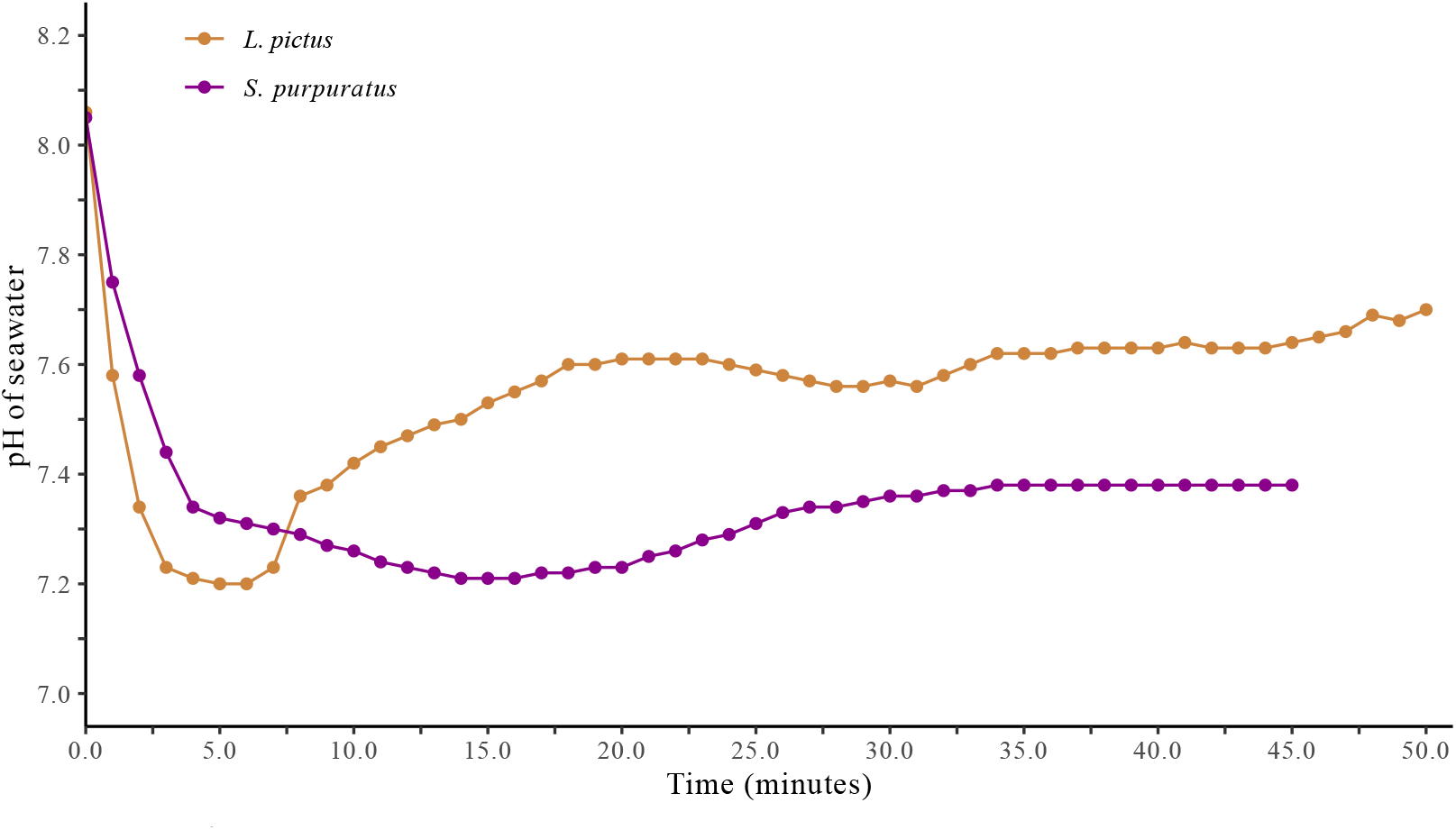
Acid release from *L. pictus* and *S. purpuratus* sperm upon dilution into seawater. One hundred microliters of semen of each species was diluted into 5 ml FSW and the pH of the seawater measured at 1 min intervals with a combination electrode (16).

Developing cryopreservation methods for sperm are difficult because they involve so many variables. People who have not worked with sea urchin gametes cannot appreciate interspecies and intraspecies differences. For example, the sperm of one species of New Zealand sea urchin can fertilize eggs after being frozen and thawed without a cryoprotectant (1). Optimal DMSO concentrations for sperm of this species are 2.5-7.5% (1). In our study of sperm of these two California species, sperm do not survive LN freezing without a minimum of 5% DMSO, and at 5%, even though some sperm survive, there is substantial sperm lysis and chromatin release.

In both our species, embryos fertilized by cryopreserved sperm there is a higher percentage of morphologically abnormal individuals when compared to embryos fertilized by non-frozen sperm. Such sea urchin abnormalities of resulting from freezing have been noted by other laboratories. However, every fertilization test we have done with cryopreserved sperm of these two species has resulted in a large majority of normal embryos that become feeding larvae.

The presence of the egg jelly layer, which induces the sperm acrosome reaction, can also differ between species. When the jelly layer is completely washed off of *S.purpuratus* eggs, antibodies show that the acrosome inducer in jelly molecules is still present as part of the vitelline layer and the eggs are fertilizable with ease. However, with *L.pictus* eggs enough jelly can be washed off so that fertilization is reduced and fertilization can be increased by adding soluble jelly (35). Sea urchin egg jelly contains a fucose sulfate polymer that differs in chemical structure among species and is one of the components of species-specific induction of the sperm acrosome reaction (37,39).

### 4.3 Intraspecies differences

In *S.purpuratus*, the acrosome reaction inducer in egg jelly is a fucose sulfate polymer, which comes in two molecular structures. About 90% of females make eggs with one of the two types and 10% of females express both types. The differences between Fucan I and Fucan II are in the positions of sulfation of the fucose ring (2). *S.purpuratus* sperm also differ in their sensitivity of acrosome reaction induction as shown in one paper where controlled conditions showed that sperm of seven males can differ greatly in the amount of fucan needed to induce the acrosome reaction [35].

In *L.pictus*, the egg jelly contains 9- and 10-amino acid peptides collectively termed speract, which are chemoattractive to sperm and also help activate sperm motility. Individual females can differ by 10-fold in the amount of speract they produce (15). Egg lots from individual *L. pictus* females differ in production of eggs that can be parthenogenetically activated, some individual females never produce eggs capable of parthenogenetic development (4). Finally, we cite a study of one Japanese *sea urchin* species, *Hemicentrotus pulcherrimus*, showing that eggs from different females vary greatly in how long they survive at 4°C in antibiotic solutions (10), one egg batch surviving for 64 days [18].

## 5. Conclusions

We describe a method to keep sperm of two California sea urchin species viable for 4-9-weeks in suspension at 0°C. We also describe a method for cryopreservation of sperm and a standard assay for fertilizing ability.

## Abbreviations used

FSW: 2 μm filtered natural seawater;
KH7 extender: (FSW containing 5 mM HEPES buffer, 40 mM extra KCl, pH 7.0);
FSW/HEPES: FSW containing 1 mM HEPES, pH 8.1;
LN: liquid nitrogen;
DMSO: dimethyl sulfoxide;
pHi: intracellular pH.

## Acknowledgements

Professor Michael C. Collins is thanked for helpful discussions. Kasey L. Mitchell is thanked for technical help. Evan Tjeerdema is thanked for assistance with preparation of the figures. This work was supported by Funding from the NIH Office of Research Infrastructure Programs (034075) to AH.

